# Eicosapentaenoic Acid (EPA) Alleviates LPS-Induced Oxidative Stress via the PPARα–NF-κB Axis

**DOI:** 10.1101/2025.03.17.643636

**Authors:** Haya AlAbduljader, Halemah AlSaeed, Amenah Alrabeea, Ameenah Sulaiman, Mohammed J. A. Haider, Fahd Al-Mulla, Rasheed Ahmad, Fatema Al-Rashed

## Abstract

Metabolic-endotoxemia, characterized by the translocation of lipopolysaccharide (LPS) from Gram-negative bacteria into the bloodstream, is a key contributor to chronic low-grade inflammation associated with obesity and type 2 diabetes. This condition exacerbates metabolic disruptions by activating Toll-like receptor 4 (TLR4) on macrophages, leading to the release of pro-inflammatory cytokines and subsequent insulin resistance. Eicosapentaenoic acid (EPA/C20:5), an omega-3 polyunsaturated fatty acid, has demonstrated anti-inflammatory and antioxidative properties, but its precise mechanisms of action in mitigating LPS-induced stress remain unclear. This study investigates the pathways through which EPA/C20:5 alleviates LPS-induced oxidative stress and inflammation in macrophages. EPA pretreatment significantly reduced LPS-induced inflammatory responses, decreasing IL-1β and IL-6 expression and IL-1β secretion, and lowering the percentage of HLA-DR⁺ macrophages. EPA also attenuated ER stress, evidenced by reduced expression of ATF4, DDIT3, HSPA5/GRP78, BIP, and CHOP at both gene and protein levels. Oxidative stress was mitigated, as shown by decreased HIF1α expression, reduced ROS levels, and preservation of mitochondrial membrane potential. Importantly, EPA increased the expression of PPARα and FABP5 while inhibiting NF-κB activation independently of the TLR4-IRF5 pathway. The protective effects of EPA were abolished by PPARα inhibition with GW9662, indicating that EPA’s action is PPARα-dependent. This study highlights the modulatory role of EPA in alleviating LPS-induced oxidative stress and inflammation in macrophages through activation of the FABP5/PPARα/NF-κB axis, independently of TLR4-IRF5 signaling. These findings reveal a novel mechanism for EPA’s anti-inflammatory effects and suggest that targeting the FABP5/PPARα pathway may offer therapeutic potential for treating metabolic disorders associated with chronic inflammation.

## Introduction

Obesity and type 2 diabetes are rapidly emerging as significant global health challenges, frequently co-occurring with other metabolic disorders, including insulin resistance, metabolic syndrome, and cardiovascular disease (1, 2). A common feature across these conditions is chronic low-grade inflammation, which is partly attributed to increased levels of circulating proinflammatory cytokines. Although the precise triggers of this inflammatory state are not fully understood, translocation of lipopolysaccharide (LPS) from Gram-negative bacteria into the bloodstream has been suggested as an initiating factor (3). This process, known as metabolic endotoxemia, is often exacerbated by high-fat diets, which increase plasma LPS levels, promoting inflammation and metabolic disruptions (4). LPS mainly acts through Toll-like receptor 4 (TLR4) on immune cells, (primarily macrophages), where it triggers a cascade of inflammatory mediators such as IL-1β, IL-6, and TNFα. These mediators subsequently drive insulin resistance and metabolic dysfunction (5).

LPS-induced inflammation is known to contribute to endoplasmic reticulum (ER) stress and oxidative stress, both of which are pivotal in the pathogenesis of metabolic diseases. ER stress arises from an accumulation of misfolded proteins in the ER, initiating the unfolded protein response (UPR) to restore cellular homeostasis (6). Persistent ER stress, however, can lead to cell death through apoptosis. Similarly, oxidative stress, caused by an imbalance between reactive oxygen species (ROS) production and detoxification, damages cellular components, including lipids, proteins, and DNA, and further amplifies inflammatory signaling pathways (7, 8). Together, the interplay of ER stress, oxidative stress, and inflammation forms a vicious cycle that underpins the progression of obesity and type 2 diabetes.

Omega-3 polyunsaturated fatty acids (PUFAs), especially eicosapentaenoic acid (EPA/C20:5), have demonstrated protective effects against inflammation and oxidative stress (9). EPA, a 20-carbon chain with five double bonds, is a long-chain omega-3 fatty acid primarily found in fish oil. It enhances lipid and carbohydrate metabolism and exerts anti-inflammatory effects by interacting with several nuclear receptors. EPA directly modulates inflammation by engaging with peroxisome proliferator-activated receptors (PPARs), which regulate genes involved in lipid metabolism, energy balance, and inflammation (10).

Although fatty acids like EPA/C20:5 are recognized for their anti-inflammatory properties, the underlying mechanistic pathways remain poorly understood, particularly in the context of LPS-induced endotoxemia. In the present study, we aimed to investigate the mechanistic pathway through which EPA/C20:5 alleviates LPS-induced oxidative stress, with a focus on the PPARα– NF-κB axis. Using a cell culture model, we examined EPA’s effects on inflammation, membrane potential, and ROS levels. Our findings indicate that EPA’s anti-inflammatory action requires the activation of PPARα and FABP5 to inhibit NF-κB, and is independent of the TLR4-IRF5 pathway. This study provides insights into the potential therapeutic role of EPA in modulating cellular stress responses associated with obesity and metabolic diseases.

## Materials and methods

### Cell Culture

THP-1 human monocytic cells, obtained from the American Type Culture Collection (ATCC), were cultured in RPMI-1640 medium (Gibco, Life Technologies, Grand Island, USA). The medium was enriched with 10% fetal bovine serum (Gibco), 2 mM L-glutamine (Gibco), 1 mM sodium pyruvate, 10 mM HEPES, 100 µg/ml Normocin, and antibiotics comprising 50 U/ml penicillin and 50 µg/ml streptomycin (Gibco). The cells were maintained at 37°C in a humidified atmosphere containing 5% CO2.

### Macrophage Differentiation

Before stimulation, THP-1 monocytes were differentiated into macrophages as published previously (11). Briefly, cells were exposed to 10 ng/ml phorbol-12-myristate-13-acetate (PMA) for three days. This was followed by an additional three-day resting period in RPMI medium devoid of serum and PMA to ensure the cells were adequately primed for further experiments.

### Cell stimulation

Monocytes were seeded into 12-well plates (Costar, Corning Incorporated, Corning, NY, USA) at a density of 1 × 10⁶ cells per well, unless noted otherwise. Differentiation into macrophages followed the previously outlined protocol. The cells were then pre-incubated for 24 hours at 37°C with either 200 µM eicosapentaenoic acid (EPA/C20:5) or 24% bovine serum albumin (BSA) as a control. Post-treatment, the cells were exposed overnight to lipopolysaccharide (LPS, 10 ng/mL; L4391, Sigma Aldrich, Merck KGaA, Darmstadt, Germany) or 0.01% DMSO as a vehicle control. To investigate PPARα involvement, cells were pretreated for one hour with 32 nM GW9662 (biotechne TOCRIS 1312) before undergoing a 24-hour exposure to EPA/C20:5 and an overnight LPS challenge.

#### Preparation of BSA-Fatty Acid Complexes

Complexes of bovine serum albumin (BSA) and fatty acids were prepared using a method adapted from van Harken, Dixon, and Heimberg (12, 13). Briefly, a 24% (w/v) BSA solution was prepared by gradually dissolving fatty-acid-free BSA (Cat # A6003, Sigma) in 150 mM NaCl, adjusting the final pH to 7.4. The solution was then filtered through a 0.22 μm filter and aliquoted for storage at –20°C until use. To prepare the fatty acid/BSA solution, cis-5,8,11,14,17-Eicosapentaenoic acid (EPA, C20:5; Cat # E2011-10MG, Sigma) was added to the 24% BSA solution to a final concentration of 10 mM. This mixture was gently heated and stirred on a hot plate until an emulsion formed. The emulsion was then stored at –20°C until needed.

### Flow Cytometry Analysis

#### Staining of Cell-Surface Markers

Monocytic cells were seeded in 24-well plates at 0.5×10^6^ cells/well. Cells were transformed into macrophages and pre-treated as previously described. To detach macrophages, cells were harvested by gentle pipetting in ice-cold PBS supplemented with 0.5 mM EDTA (disodium ethylenediaminetetraacetic acid). Harvested cells were then resuspended in FACS staining buffer (BD Biosciences) and blocked with human IgG (Sigma; 20 μg) for 30 minutes on ice. Cells were washed and resuspended in 100 µl of FACS buffer and incubated with anti-CD11b PE-Cy7 (Cat # 557743; BD BD Pharmingen™) and anti-HLA-DR-PerCP 5.5 (Cat # 560652; BD Pharmingen™) or suitable isotype control antibody (Cat # 558055; BD Phosflow™, Cat # 559529; BD Phosflow™ , Cat # 560542; BD Pharmingen™ or Cat # 560817; BD Phosflow™) on ice for 30 minutes.

Cells were washed three times in FACS buffer to remove nonspecific binding then resuspended in FACS buffer for FACS analysis (FACSCanto ; BD Bioscience, San Jose, USA). FACS data analysis was performed using BD FACSDiva^TM^ Software 8 (BD Biosciences, San Jose, USA) or FlowJo. Unstained cells were used to set the quadrant of the negative *vs* positive gates, and the Median Fluorescence Intensity (MFI) was measured. Gating strategy is summarized in **Supplementary Figure 1A**.

#### Staining of intracellular markers

For intracellular staining, cells were incubated with fixation/permeabilization buffer (Cat # 00-5523-00, eBioscience, San Diego, CA, USA) for 20 min at 4°C, followed by washing with Primwash solution. Cells were then stained with anti-hIRF5 Alexa Flour ® 488 (Cat # IC4508G; R&D systems) and anti-NF-κB p65 (pS529)-PE (Cat # 558423; BD Bioscience). Samples were processed in BD FACSCanto and FACS data analysis was performed using BD FACSDiva^TM^ Software 8 (BD Biosciences, San Jose, USA) or FlowJo. Unstained cells were used to set the quadrant of the negative *vs* positive gates, and the Median Fluorescence Intensity (MFI) was measured. Gating strategy is summarized in **Supplementary Figure 1B**.

#### Analysis of Intracellular Reactive Oxygen Species (ROS) by flow cytometry

To assess intracellular ROS levels, we used the cell-permeable fluorogenic probe 2’,7’-Dichlorodihydrofluorescin diacetate (DCFH-DA / Bioquochem ; KP06003). This probe enters cells and is converted by intracellular nonspecific esterases into highly fluorescent 2’,7’-dichlorofluorescein (DCF) upon reaction with ROS, serving as an indicator of intracellular oxidative stress. The assay was performed following the manufacturer’s instructions. Briefly, THP-1 transformed macrophages were preincubated with 10 µM DCFH-DA at 37°C for 30 minutes. Cultures were then pre-treated with EPA (200 µM) and/or GW9662 (PPARα antagonist) or positive control (50 μM tert-Butyl hydroperoxide (TBHP) solution) followed by LPS (10 ng/ml) stimulation. After 4 hours of incubation, cells were harvested, rinsed three times with PBS. DCF fluorescence was measured by flow cytometry with excitation and emission wavelengths of 488 nm and 525 nm, respectively. For each sample, 10,000 events were recorded. Data analysis was conducted to quantify ROS levels as a measure of oxidative stress in different treatment groups.

### Measurement of Mitochondrial Membrane Potential with JC-1

Changes in mitochondrial membrane potential were assessed using JC-1 (5,5’,6,6’-tetrachloro-1,1’,3,3’-tetraethyl-benzimidazolcarbocyanine iodide, Molecular Probes) (cat.#: T3168; Thermofisher). JC-1 emits either green or red fluorescence: a green signal indicates depolarized (low membrane potential) mitochondria, while a red signal indicates polarized (high membrane potential) mitochondria (14). THP-1 transformed macrophages were pre-treated with EPA (200 µM) and/or GW9662 (PPARα antagonist) followed by stimulation with LPS (10 ng/ml) as per experimental design. After 24 hours of incubation, cells were harvested by ice cold PBS supplemented with 0.5 mM EDTA (Ethylenediaminetetraacetic acid, disodium salt). Cells were centrifugated at 300 x g for 5 minutes and washed twice with pre-warmed PBS. Cells were then resuspended in 2 µM JC-1 and incubated at 37°C for 20 minutes in the dark. Following incubation, cells were washed twice with PBS to remove excess dye and resuspended in 500 ml PBS for analysis.

JC-1 fluorescence was measured using FACSCanto II with laser excitation at 488 nm. Emissions were detected at two channels: green fluorescence for JC-1 monomers (depolarized mitochondria) at 530/30 nm and red fluorescence for JC-1 aggregates (polarized mitochondria) at 585/42 nm. For each sample, at least 10,000 events were recorded. Data analysis was performed using FlowJo software, and the mitochondrial membrane potential was determined by calculating the red-to-green fluorescence ratio. A higher ratio reflects preserved mitochondrial membrane potential, while a lower ratio indicates mitochondrial depolarization.

### Quantitative Real-Time Polymerase Chain Reaction (qRT-PCR)

Total RNA was isolated using the RNeasy Mini Kit (Qiagen, Valencia, CA, USA) following the manufacturer’s protocol. Complementary DNA (cDNA) was synthesized from 1 μg of RNA using the High-Capacity cDNA Reverse Transcription Kit (Applied Biosystems, Foster City, CA, USA). Quantitative real-time PCR (qRT-PCR) was carried out on the 7500 Fast Real-Time PCR System (Applied Biosystems) using the TaqMan® Gene Expression Master Mix (Applied Biosystems). Each reaction utilized 1000 ng of cDNA and TaqMan Gene Expression Assay reagents, as detailed in Table 1. Threshold cycle (Ct) values were normalized to GAPDH, and the ΔΔCt method was employed to calculate relative mRNA levels compared to the control. Relative gene expression was presented as fold change, with the control set to 1. Data are shown as mean ± SEM, and statistical analysis was conducted with significance determined at p < 0.05.

**Table 1:** List of qRT-PCR Primers.

| REAGENT OR RESOURCES | SOURCE | IDENTIFIER |
| --- | --- | --- |
| **IL-1B (HUMAN)** | ThermoFisher | Hs01555410_m1 |
| **IL-6 (HUMAN)** | ThermoFisher | Hs00174131_m1 |
| **TNFA(HUMAN)** | ThermoFisher | Hs01113624_g1 |
| **ATF4 (HUMAN)** | ThermoFisher | Hs00909569_g1 |
| **DDIT3 (HUMAN)** | ThermoFisher | Hs00358796_g1 |
| **HSPA5(HUMAN)** | ThermoFisher | Hs00607129_gH |
| **PPARA(HUMAN)** | ThermoFisher | Hs00947536_m1 |
| **PPARD(HUMAN)** | ThermoFisher | Hs04187066_g1 |
| **PPARG (HUMAN)** | ThermoFisher | Hs01115513_m 1 |
| **HIF1A (HUMAN)** | ThermoFisher | Hs00153153_m1 |
| **FABP4 (HUMAN)** | ThermoFisher | Hs01086177_m1 |
| **FABP5 (HUMAN)** | ThermoFisher | Hs02339439_g1 |
| **IRF4 (HUMAN)** | ThermoFisher | Hs00180031_m1 |
| **IRF5 (HUMAN)** | ThermoFisher | Hs00158114_m1 |
| **GAPDH (HUMAN) (VIC-**<br>**TAMARA)** | Applied biosystems<br>ThermoFisher | 4310859 |

### Western Blot Analysis of Protein Expression

Treated cell cultures were lysed using RIPA buffer (Thermo Scientific) supplemented with a protease and phosphatase inhibitor cocktail (Thermo Fisher Scientific) to safeguard against protein degradation. The lysates were centrifuged at 14,000 x g for 15 minutes at 4°C to eliminate debris. Protein concentration was determined using the Bradford assay (Bio-Rad), and equal amounts of protein (12.5 µg per sample) were resolved on a 10% SDS-PAGE gel by electrophoresis. Subsequently, proteins were transferred to a PVDF membrane (Millipore) via a wet transfer method.

To block non-specific binding, the membrane was incubated at room temperature for 1 hour in 5% non-fat dry milk dissolved in Tris-buffered saline with 0.1% Tween-20 (TBST). After blocking, the membrane was treated overnight at 4°C with primary antibodies specific to the proteins of interest (as listed in Table 2), with β-Actin serving as a loading control. The following day, membranes were washed with TBST and incubated with the appropriate horseradish peroxidase (HRP)-conjugated secondary antibody for 1 hour at room temperature.

**Table 2:** List of Western Blot Primary Antibodies.

| NAME (WESTERN<br>BLOT) | SPECIES | COMPANY | REFERENCE/CATALOG<br>NUMBER |
| --- | --- | --- | --- |
| **BIP (C50B12)** | Rabbit mAb | Cell signalling<br>technology | 3177 |
| **CHOP (L63F7)** | Mouse mAb | Cell signalling<br>technology | 2895S |
| **BETA-ACTIN**<br>**(8H10D10)** | Mouse mAb | Cell signalling<br>technology | 3700S |
| **PPAR ALPHA**<br>**(PHOSPHO S12)** | Rabbit pAb | abcam | ab3484 |
| **FABP4 (D25B3) (XP) ®** | Rabbit mAb | Cell signalling<br>technology | 3544S |
| **FABP5 (D1A7T)** | Rabbit mAb | Cell signalling<br>technology | 39926S |

Protein bands were visualized using an enhanced chemiluminescence (ECL) reagent (Thermo Scientific) and documented using a gel imaging system (Bio-Rad). Quantification of protein expression was conducted using ImageJ software, with normalization of target protein levels to β-Actin for consistency.

### Measurement of AP-1/NF-κB activity

THP-1 XBlue monocytes (InvivoGen, San Diego, CA) are engineered to stably express a reporter construct containing the secreted embryonic alkaline phosphatase (SEAP) gene, which is regulated by a promoter responsive to the transcription factors AP-1 and NF-κB. Activation of NF-κB triggers increased SEAP secretion. Cells were pretreated following the previously described protocol(15). SEAP levels were measured in the culture supernatants after 3-4 hours of incubation with Quanti-Blue solution (InvivoGen) at 650 nm using an ELISA plate reader.

### Sandwich Enzyme-Linked Immunosorbent Assay (ELISA)

Secreted IL-1β protein concentrations were quantified in the media of THP-1 treated cell cultures using sandwich ELISA in accordance with the manufacturer’s instructions (R&D systems, Minneapolis, MN, USA , Cat # DY279) and as previously published (16) .

### Statistical Analysis

Statistical analysis was performed using GraphPad Prism software (La Jolla, CA, USA). Data are shown as the mean ± the standard error of the mean (SEM), unless otherwise indicated. Parametric data were analyzed by one-way ANOVA followed by Tukey’s post hoc multiple comparisons test. For all analyses, data from a minimum of three replicates were used for statistical calculation. A *p* value of 0.05 was considered statistically significant. ns = non-significant, * *p* < 0.05, ** *p* < 0.01, *** *p* < 0.001, and **** *p* < 0.0001.

## Results

### EPA downregulates LPS-induced inflammatory response and ER stress

The anti-inflammatory properties of omega-3 fatty acids have been well-documented in numerous studies. However, despite the variety of omega-3 family members, their specific effects on inflammatory outcomes remain partially unclear, with some demonstrating more potent effects in mitigating inflammation than others. One proposed mechanism is that these fatty acids alleviate endoplasmic reticulum (ER) stress, thereby reducing the inflammatory response.

To test this hypothesis and further understand the underlying mechanisms, we utilized a THP-1 derived macrophage model. The cells were pre-incubated with 200 µM of EPA/C20:5 and subsequently stimulated with 10 ng/ml of LPS. As expected, pre-treatment with EPA significantly reduced the percentage of HLA-DR^+^ macrophage subsets compared to cells exposed only to LPS (**Figure 1A and B**) verifying this anti-inflammatory property.

**Figure 1:**
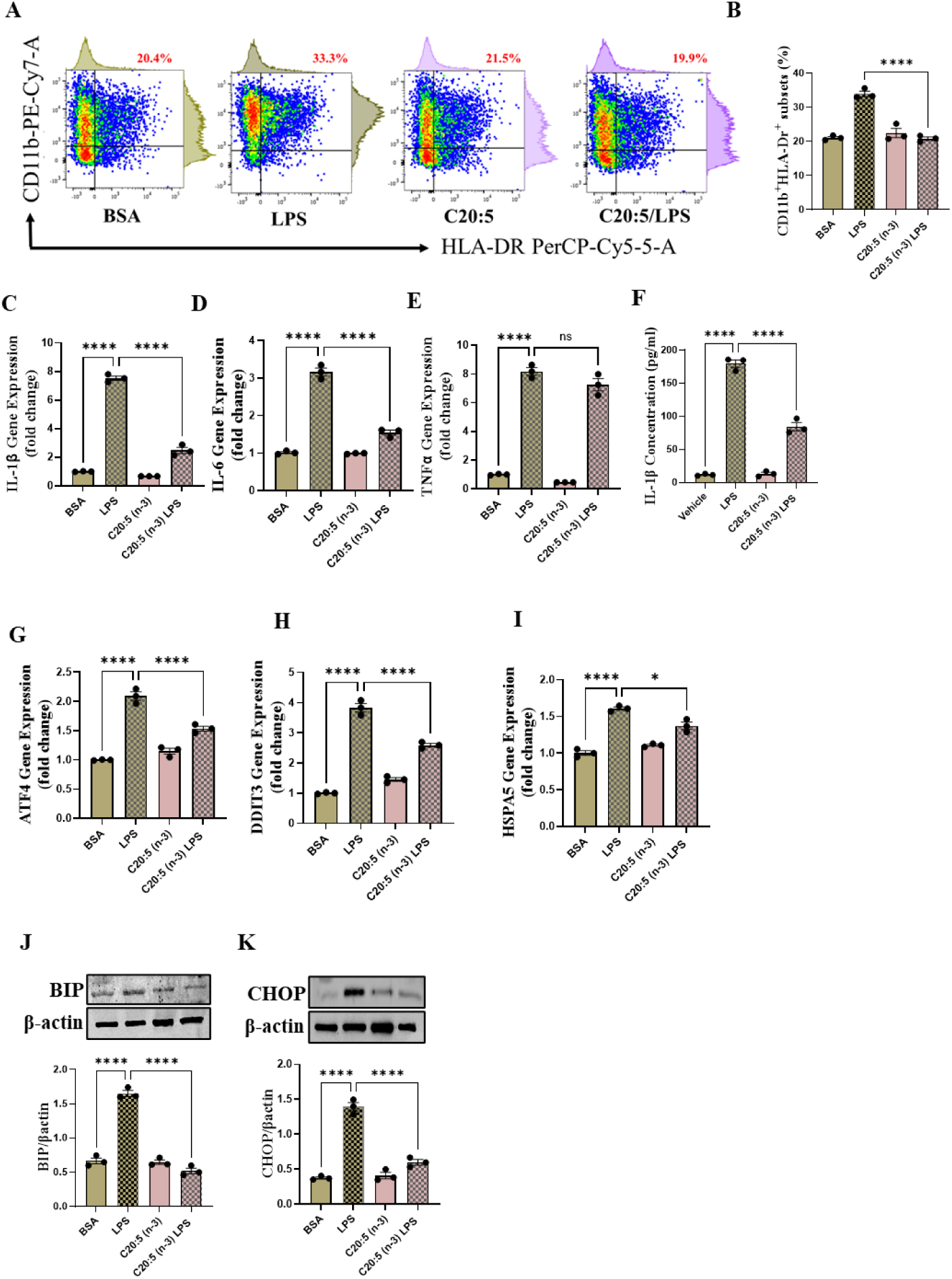
Eicosapentaenoic Acid (EPA/C20:5) Reduces LPS-Induced Inflammatory Response, ER Stress, and Oxidative Stress in THP-1 derived Macrophages. THP-1 macrophages were pre-treated with EPA (200 µM) and subsequently stimulated with LPS (10 ng/ml). (A) Representative dot plots of flow cytometry analysis of HLA-DR^+^ macrophage subsets. (B) Bar graph analysis of HLA-DR^+^ subsets percentage. Quantitative gene expression analysis of (C) IL-1β, (D) IL-6, and (E) TNF-α in THP-1 macrophages. (F) Protein secretion profile of IL-1β in the culture supernatant. Quantitative gene expression analysis of ER stress markers (G) ATF4, (H) DDIT3, and (I) HSPA5 (GRP78) in response to EPA/C20:5 treatment. Western blot analysis of (J) BIP and (K) CHOP protein levels corrected to β-actin. Data are presented as mean ± SEM, with a minimum of *n* = 3. Statistical significance was assessed using one-way ANOVA followed by Tukey’s post-hoc test. \**p* < 0.05, and \*\*\*\**p* < 0.0001.

This effect was also observed at the gene level, where EPA pre-treatment significantly downregulated the gene expression of pro-inflammatory cytokines IL-1β and IL-6 (**Figure 1C and D**). Although a reduction in TNF-α gene expression was observed, it did not reach statistical significance (**Figure 1E**). Furthermore, we tested the secretion profile of these pro-inflammatory cytokines in the media. As expected, a significant reduction was seen in IL-1β in cells that were pretreated with EPA/C20:5 (**Figure 1F**). These results indicate that EPA effectively mitigates LPS-induced inflammatory responses.

Previous studies have demonstrated that LPS induces ER stress (17). In our model, EPA/C20:5 pre-treatment significantly reduced the expression of key ER stress markers, including ATF4, DDIT3, and HPAS5/GRP78 at the gene level (**Figure 1G-I**). This reduction in ER stress was further confirmed at the protein level, as evidenced by decreased expression of BIP, a chaperone associated with HPAS5, and CHOP, a protein marker for DDIT3 (**Figure 1J and K**). Altogether, these findings suggest that EPA/C20:5 attenuates both the inflammatory response and ER stress induced by LPS.

### EPA downregulates LPS-induced oxidative stress

To further explore the mechanisms by which EPA mitigates cellular stress, we examined its impact on oxidative stress, a well-known contributor to ER stress and inflammation. Given the role of HIF1α in promoting oxidative stress during inflammatory conditions, we first assessed its gene expression, suggesting that EPA/C22:5 might mitigate oxidative stress through modulation of this pathway. As expected, LPS stimulation showed significant upregulation in HIF1α gene expression, indicating an induction of oxidative stress similar to our previous observations where pre-treatment with EPA/C20:5 prior to LPS stimulation significantly reduced this upregulation (**Figure 2A**). To further verify this observation, we conducted a functional analysis for ROS production using a flow cytometric analysis of DCFH-DA. Consistent with a reduction in HIF1α expression, ROS levels were markedly reduced following EPA/C20:5 treatment (**Figure 2B**). This indicates that EPA effectively alleviates oxidative stress, potentially through HIF1α downregulation and related pathways. Furthermore, since mitochondrial dysfunction is a major source of ROS production, we further examined the effect of EPA/C20:5 on mitochondrial membrane potential using a JC-1 assay. Compared with the LPS-treated group, pre-treatment with EPA/C20:5 preserved mitochondrial membrane potential as indicated by a stable aggregate to monomers ratio (**Figure 2C and D**), suggesting that EPA/C20:5 prevents mitochondrial depolarization under LPS-induced stress conditions. Collectively, the observed data underscore the multifaceted protective role of EPA/C20:5 in cellular stress responses, likely mediated through its impact on mitochondrial integrity and oxidative stress regulation.

**Figure 2:**
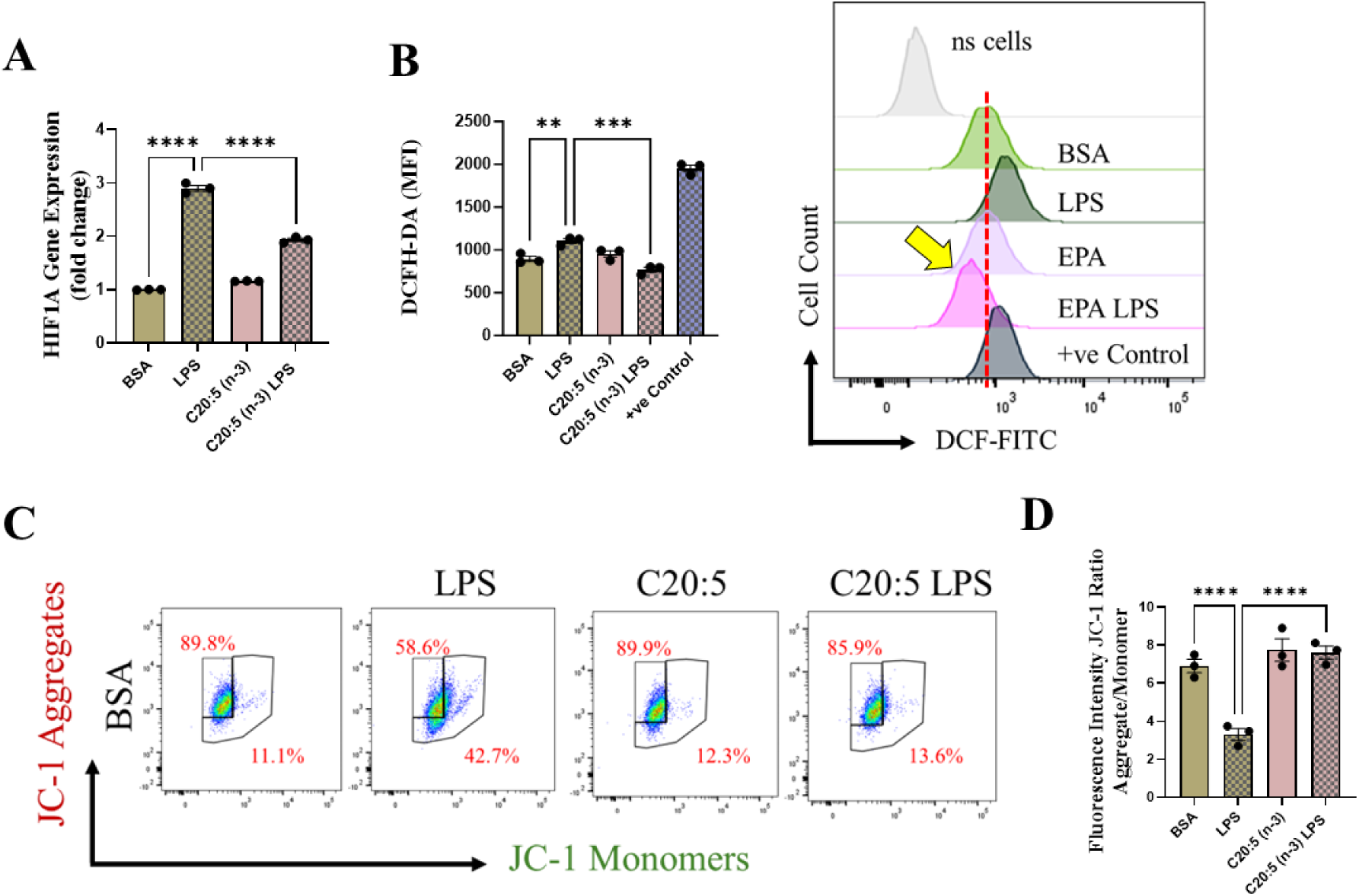
Eicosapentaenoic Acid (EPA) Preserves Mitochondrial Membrane Potential and Reduces Oxidative Stress in LPS-Stimulated THP-1 derived Macrophages. (A) Quantitative gene expression analysis of HIF1A. (B) Intracellular ROS was measured by flow cytometry (DCFH-DA) with data presented as a bar graph of median fluorescence intensity (MFI) with a representative histogram. (C) Representative dot plots from flow cytometry for JC-1 aggregates (representing polarized mitochondria) and monomers (representing depolarized mitochondria). (D) Bar graphs show aggregate-to-monomer ratio. Data are presented as mean ± SEM, with a minimum of *n* = 3. Statistical significance was determined using one-way ANOVA followed by Tukey’s post-hoc test. \*\**p* < 0.01, \*\*\**p* < 0.001, and \*\*\*\**p* < 0.0001.

### EPA modulates FABP5/PPARα/NF-κβ signaling independently of the TLR4-IRF5 pathway

To further elucidate the molecular mechanisms underlying the protective effects of EPA/C20:5, we focused on key modulators involved in lipid metabolism and inflammation. Specifically, we investigated the expression of PPARα, IRF5, and FABP4/5 given their role in mitochondrial function, oxidative stress, and inflammation. PPARα, a nuclear receptor that regulates mitochondrial fatty acid oxidation and energy homeostasis, was significantly reduced by LPS treatment. Interestingly, EPA/C20:5 elevated PPARα expression at both gene and protein levels (**Figure 3A and B, respectively**). This observation suggests a key role in re-establishing mitochondrial function and metabolic homeostasis in the presence of inflammatory stimuli such as LPS. Fatty acids (FA), including EPA/C20:5, are known ligands for PPARs. Their subsequent binding can impact their activity. During this process, FA requires the initial interaction with fatty acid binding proteins (FABP), which act as chaperones to transport ligands to PPARs. While no significant effect was observed on FABP4 expression when compared to LPS-treated cells, EPA/C20:5 induced a marked increase in FABP5 at both gene and protein levels **(Figure 3C-E**). This observation indicates that EPA/C20:5 might exert its protective effects through this specific pathway, facilitating its interaction with PPARα to promote fatty acid oxidation and anti-inflammatory responses. Given that IRF5 is a key transcription factor activated downstream of Toll-like receptor 4 (TLR4) signaling, we explored its role in modulating the inflammatory response through the activation of the NF-κβ pathway. At the gene level, the expression of IRF5 was abolished under EPA/C20:5 pre-treatment (**Figure 3F**). Subsequently, EPA/C20:5 pre-treatment significantly reduced NF-κβ activation as demonstrated in THP-1 -NF-κβ reporter cells, where we observed a notable reduction in NF-κβ expression (**Figure 3G**). To further understand this dynamic and to validate the inhibitory effect of EPA/C20:5 on the IRF5/NF-κβ pathway, we conducted flow cytometric analysis. Interestingly, while there was no significant change in the percentage of IRF5^+^ subsets, we observed a significant reduction in NF-κβ MFI within these cells, indicating that EPA’s primary mode of action may not directly inhibit IRF5 expression but instead suppress NF-κβ activation downstream of the TLR4-IRF5 axis (**Figure 3H**). Together, these results highlight the capacity of EPA/C20:5 to modulate NFκβ through the FABP5/PPARα/ NF-κβ axis independently of TLR4/IRF5.

**Figure 3:**
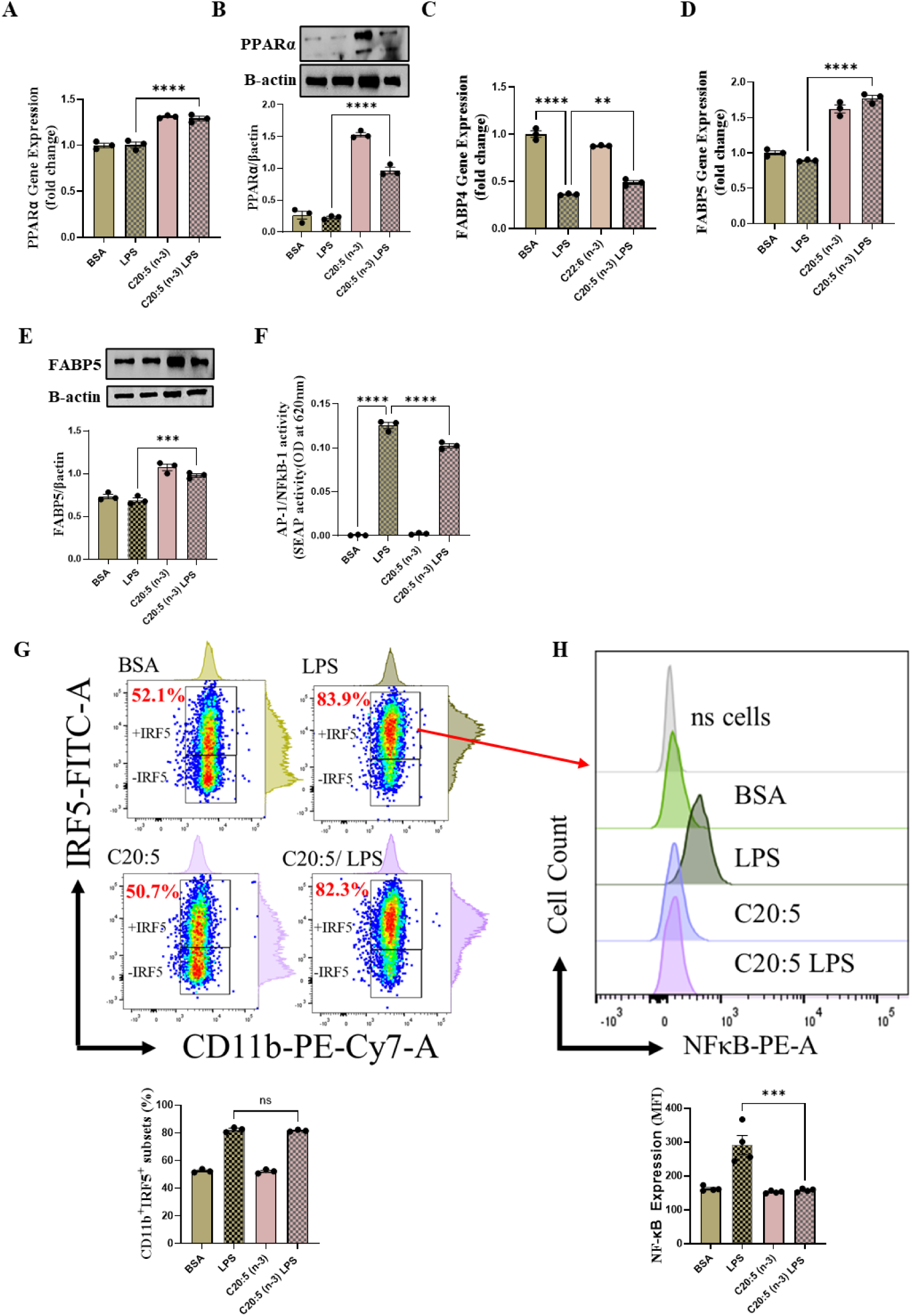
EPA/C20:5 Modulates the PPARα/ FABP5/ NF-κB pathway independently of IRF5 Activation. Quantitative analysis of PPARα expression at the (A) mRNA and (B) protein levels. (C) Quantitative analysis of FABP4 gene expression. Quantitative analysis of FABP5 expression at the (D) mRNA and (E) protein levels. (F) NF-κB reporter monocytic cells SEAP reporter activity. (G) Representative dot plots show the percentage of CD11b^+^IRF5^+^ subsets and bar graph showing the percentage. (H) Representative flow cytometry histogram and bar graph depicting the median fluorescence intensity (MFI) of NF-κB in CD11b^+^IRF5^+^ subsets. Data are presented as mean ± SEM, with a minimum of *n* = 3. Statistical significance was determined using one-way ANOVA followed by Tukey’s post-hoc test. \*\**p* < 0.01, \*\*\**p* < 0.001 and \*\*\*\**p* < 0.0001.

### PPARα inhibition abrogates EPAs effect on LPS stress and inflammation

To further delineate the role of PPARα in mediating the protective effects of EPA, we pretreated macrophages with GW9662, a potent PPARα antagonist, prior to EPA/C20:5 treatment and LPS stimulation. Under PPARα inhibition, it became evident that EPA’s ability to mitigate oxidative stress and subsequent ER stress was significantly diminished, as observed in the expression of HIF1α (**Figure 4A**). This loss of function was also noted in ER stress genes, including DDIT3, ATF4, and HSPA5, which were elevated to levels comparable to those seen with LPS stimulation alone (**Figures 4B-D**). This outcome was further substantiated by the expression of BIP protein, a critical marker of ER stress (**Figure 4E**), and an elevation of the pro-inflammatory marker IL-1β in the media, with no significant impact observed in EPA/C20:5-treated cells under PPARα inhibition (**Figure 4F**). Additionally, both functional assays for ROS production (**Figure 4G**) and mitochondrial depolarization (**Figures 4H and I**) showed no significant protection in EPA/C20:5-treated cells under PPARα inhibition, as the differences between EPA/C20:5-treated and untreated groups were not statistically significant. These observations indicate that the protective effects of EPA/C20:5 were nullified by PPARα inhibition.

**Figure 4:**
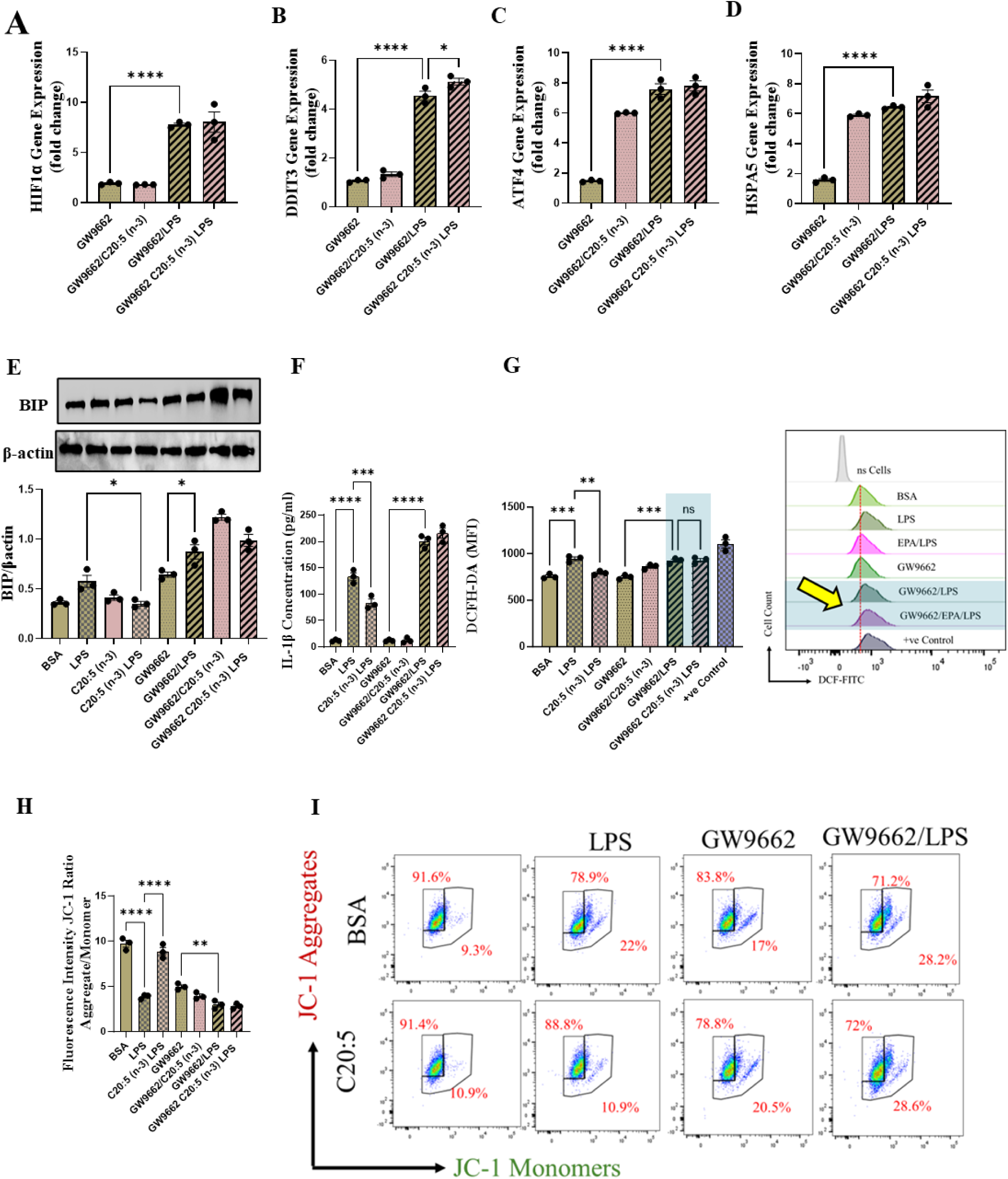
PPARα Antagonism Abrogates EPA’s Protective Effects on LPS-Induced Inflammatory and Cellular Stress Responses in THP-1 derived Macrophages. Cells were pre-treated with EPA (C20:5, 200 µM), GW9662 (a PPARα antagonist), or both, followed by LPS (10 ng/ml) stimulation. Quantitative analysis of gene expression for ER stress markers (A) HIF1A, (B) DDIT3 , (C) ATF4, and (D) HSPA5. (E) Western blot analysis of BIP protein level. (F) Protein secretion analysis of IL-1β (G) Bar graph and representative histogram of ROS production measured by DCFH-DA assay. (H) Quantification of mitochondrial membrane potential using a JC-1 assay. (I) Flow cytometric dot plots for JC-1 aggregates and monomers showing mitochondrial polarization status across different treatment groups. Data are presented as mean ± SEM, with a minimum of *n* = 3. Statistical significance was determined using one-way ANOVA followed by Tukey’s post-hoc test. Significance levels are indicated as follows: \**p* < 0.05, \*\**p* < 0.01, \*\*\**p* < 0.001, \*\*\**p* < 0.0001; ns = not significant.

Collectively, these findings demonstrate that PPARα activation is essential for EPA/C20:5 to exert its protective effects against LPS-induced oxidative and ER stress. The loss of EPA/C20:5’s beneficial effects in the presence of PPARα inhibition highlights the critical role of PPARα in modulating cellular responses to inflammatory and stress stimuli.

## Discussion

Our study demonstrates that EPA/C20:5 alleviates LPS-induced oxidative stress and inflammation through a mechanism involving the PPARα–NF-κB axis, with a pivotal role for FABP5 as a chaperone in facilitating EPA’s binding to PPARα. These findings contribute to a growing body of research on the anti-inflammatory and cytoprotective effects of PUFAs in inflammatory and metabolic disorders, offering new insights into the precise molecular pathways involved.

The association between high-fat diets, increased circulating LPS levels, and chronic low-grade inflammation is well-established, with elevated LPS levels contributing to the development of obesity and insulin resistance through TLR4-mediated signaling pathways (8, 18). LPS-induced inflammation not only promotes insulin resistance but also induces ER stress and oxidative stress, both of which play critical roles in the pathogenesis of metabolic diseases (19, 20). ER stress arises from the accumulation of misfolded proteins, activating the unfolded protein response (UPR), which, when unresolved, can lead to apoptosis (21). In our model, EPA pre-treatment significantly reduced the expression of ER stress markers, such as ATF4, DDIT3, and HSPA5, suggesting that EPA can attenuate LPS-induced ER stress and restore cellular homeostasis. These findings align with previous studies highlighting omega-3 PUFAs’ ability to reduce ER stress, which is known to contribute to the progression of obesity-related metabolic inflammation (22, 23).

Oxidative stress is another key factor exacerbating inflammation in metabolic disorders, as excessive ROS production can damage cellular components and activate inflammatory signaling pathways, including NF-κB (24). In this study, EPA/C20:5 significantly reduced ROS levels and preserved mitochondrial membrane potential, as shown by a stable aggregates-to-monomers ratio in the JC-1 assay, indicating that EPA prevents mitochondrial dysfunction, a major source of ROS production during inflammatory conditions (25). Our findings align with the work of An *et al*., who demonstrated that long-term omega-3 fatty acids supplementation mitigated tubulointerstitial injury in animals with chronic renal disease by reducing oxidative stress, inflammation, and fibrosis (26). Furthermore, we observed a reduction in HIF1α expression following EPA treatment, which likely contributes to the decrease in ROS production. HIF1α is known to be upregulated during hypoxic and inflammatory conditions, promoting oxidative stress (27). EPA’s downregulation of HIF1α suggests a potential pathway by which it mitigates oxidative stress and maintains cellular integrity under inflammatory stress.

One interesting observation in our study is the essential role of PPARα and FABP5 in mediating EPA’s protective effects. PPARα, a nuclear receptor involved in lipid metabolism and anti-inflammatory responses, has been previously associated with the beneficial effects of omega-3 fatty acids in metabolic health (28). However, our study highlights a specific role for PPARα in modulating the inflammatory response to LPS, which has not been widely explored. We demonstrated that EPA/C20:5 increases PPARα expression and requires its activation to exert anti-inflammatory and cytoprotective effects. This finding is supported by studies showing that PPARα activation is crucial in reducing ER stress and inflammation in obesity models (29). The addition of FABP5 as a key mediator in this pathway provides further insight, in that FABP5 serves as a chaperone for long-chain fatty acids, facilitating their transport to PPARs (30). The observed increase in FABP5 expression following EPA treatment suggests that EPA relies on this chaperone’s function to effectively activate PPARα, thereby enhancing its anti-inflammatory properties.

Interestingly, our data show that EPA’s effects on NF-κB are independent of the TLR4-IRF5 pathway. Although IRF5 is an essential downstream component of TLR4 signaling in the activation of NF-κB (31), EPA’s reduction of NF-κB activation occurred without significant alteration in IRF5^+^ subsets. This suggests that EPA/C20:5 might inhibit NF-κB activation through a pathway distinct from canonical TLR4-IRF5 signaling. Recent studies have indicated that PPARα can directly interfere with NF-κB activity, offering an alternative route for EPA’s anti-inflammatory effects (32). Our findings contribute to this understanding by demonstrating that EPA’s anti-inflammatory action can bypass TLR4 signaling, instead relying on FABP5/PPARα to inhibit NF-κB.

To further confirm the role of PPARα in EPA’s protective effects, we employed GW9662, a selective PPARα antagonist, to inhibit PPARα activity. Under PPARα inhibition, EPA’s effects on ER stress, oxidative stress, and cytokine production were significantly reduced, indicating that PPARα activation is indeed necessary for EPA’s anti-inflammatory and cytoprotective actions. Additionally, the functional assays revealed that EPA’s ability to reduce ROS production and maintain mitochondrial membrane potential was abolished with PPARα inhibition, further confirming the receptor’s role in regulating mitochondrial and oxidative stress responses.

Overall, our study provides a mechanistic framework for EPA’s protective effects in LPS-induced cellular stress, emphasizing the roles of PPARα and FABP5 in modulating the NF-κB pathway independently of TLR4-IRF5 signaling (Figure 5).

**Figure 5:**
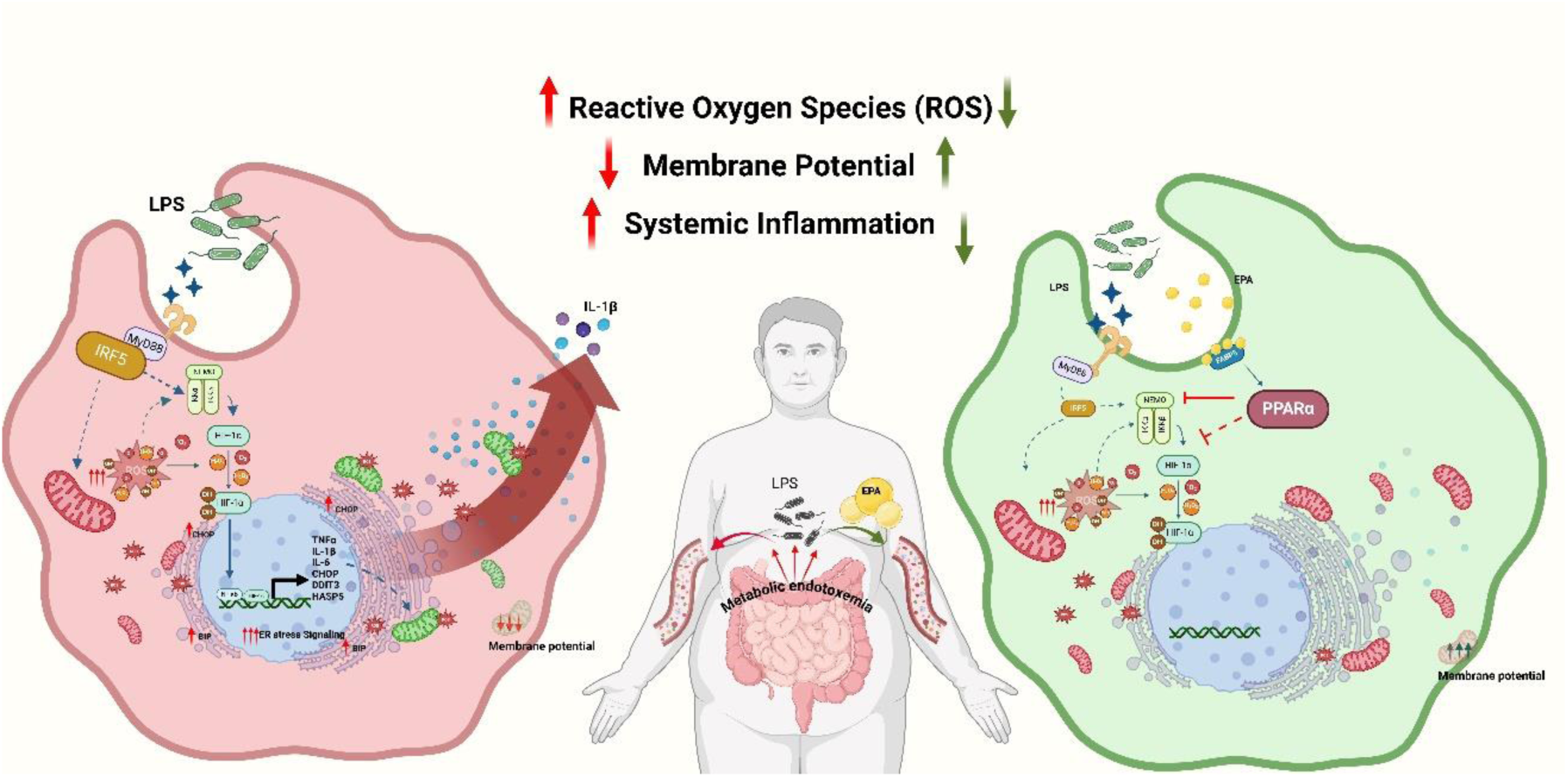
schematic representation of the study findings. Metabolic endotoxemia as a key driver of systemic inflammation through the up regulation of LPS in circulation. The left panel illustrates the impact of LPS stimulation on macrophages, leading to increased production of reactive oxygen species (ROS), loss of mitochondrial membrane potential, and activation of inflammatory pathways mediated by HIF-1α, NF-κB, and ER stress markers (CHOP, DDIT3, and HSPA5). This results in heightened systemic inflammation. The right panel demonstrates the protective effects of EPA treatment on LPS-stimulated macrophages. EPA enhances mitochondrial membrane potential, reduces ROS, and downregulates inflammatory markers, partly through the activation of FABP5/PPARα, which mitigates HIF-1α , NF-κB and ER stress signaling, ultimately alleviating systemic inflammation. This schematic underscores the therapeutic potential of EPA in reducing inflammation and oxidative stress associated with endotoxemia

These findings suggest that targeting the FABP5/PPARα axis may offer a therapeutic strategy for mitigating inflammation and cellular stress linked to obesity and metabolic disorders. Given the demonstrated benefits of dietary EPA intake in reducing cardiovascular and metabolic risks, future studies should further investigate its therapeutic potential, alongside other PPARα agonists, in clinical models of metabolic endotoxemia. Exploring interactions between EPA and cellular stress pathways may also reveal novel approaches for managing chronic inflammation and oxidative stress in metabolic diseases.

## Contributors

HAA. conducted experiments, data curation, analyzed data, interpreted the results, and participated in writing the original draft of the manuscript. HAS. And AS participated in conducting experiments, data collection, data analysis, data interpretation and wrote original draft of the manuscript. AAL and MJH participated in study designing, data interpretation, wrote original draft of the manuscript, review & editing manuscript. FAM. participated in the study designing, data interpretation, review & editing manuscript. RA. conceived idea, participated in study designing, interpreted the results, acquired funds, reviewed & editing manuscript. FAR. conceived idea, acquired funds, designed study, interpreted the results, verified data, wrote the manuscript. All authors contributed to reviewing the paper and all authors have read and approved the final version for submission.

## Conflict of Interest

The authors declare no conflicts of interest regarding the publication of this manuscript.

## Data Availability

The data supporting the findings of this study are available from the corresponding author upon reasonable request.

## Acknowledgements

This work was supported by Kuwait Foundation for the Advancement of Sciences (KFAS) Grant RA CB-2019-002 (FAR.). The funders had no role in the study design; collection, analysis and interpretation of data; writing of this paper; or in the decision to submit the paper for publication.

## Supplementary Material

The supplementary material includes the gating strategy for the flow cytometry data and raw Western blot immunoblot images, providing additional methodological details and unprocessed data to support the findings presented in the manuscript.

## Notes

### Competing Interest Statement

The authors have declared no competing interest.

