## Supplementary Figure 1 for "Eicosapentaenoic Acid (EPA) Alleviates LPS-Induced Oxidative Stress via the PPARα–NF-κB Axis"

### Supplementary Figure S1

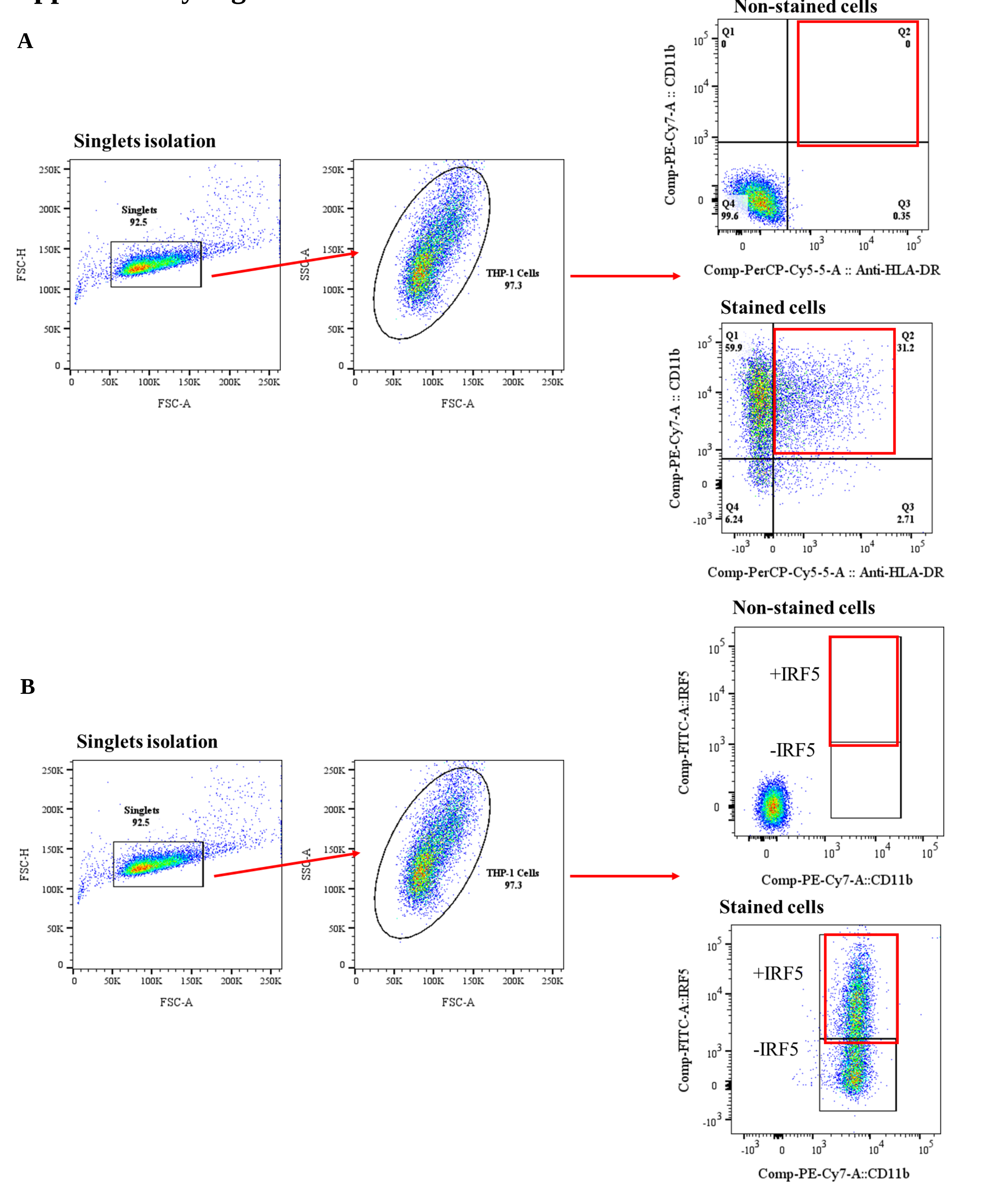


Figure S1: The gating strategy used for flow cytometric analysis in THP-1 cells is shown. Initially, a forward scatter (FSC) area versus FSC height plot was used to isolate single cells, thereby excluding doublets and cell aggregates. This "Singlets" gate, ensuring that only single-cell events were included in subsequent analysis. Next, a side scatter (SSC) versus FSC plot was applied to the singlet population to identify the THP-1 macrophage population based on cell size and granularity. Cells within this gate, labeled "THP-1 Cells," represented 92-98% of the singlet population, providing a highly purified subset of THP-1 macrophages for further examination. (A) To identify the pro-inflammatory profile, THP-1 cells were gated on a CD11b versus HLA-DR plot, selecting for CD11b+HLA-DR+ subsets. (B) For the isolation and identification of IRF5+ subsets, cells were gated on a CD11b versus IRF5 plot to identify the CD11b+IRF5+ population. This sequential gating approach ensures precise selection of specific cell subsets for downstream analysis.
